# Strong and specific associations between cardiovascular risk factors and brain white matter micro- and macro-structure in health ageing

**DOI:** 10.1101/264770

**Authors:** Delia Fuhrmann, David Nesbitt, Meredith Shafto, James B. Rowe, Darren Price, Andrew Gadie, Cam-CAN, Rogier A. Kievit

## Abstract

Cardiovascular health declines with age, increasing the risk of hypertension and elevated heart rate in middle- and old age. Here, we used multivariate techniques to investigate the associations between cardiovascular health (diastolic blood pressure, systolic blood pressure and heart rate) and white matter macrostructure (lesion volume and number), and microstructure (as measured by Diffusion Weighted Imaging) in the cross-sectional, population-based Cam-CAN cohort (*N* = 667, aged 18 to 88). We found that cardiovascular health and age made approximately similar contributions to white matter health and explained up to 56% of variance. Lower diastolic blood pressure, higher systolic blood pressure and higher heart rate were each strongly, and independently, associated with white matter abnormalities on all indices. Body mass and exercise were associated with white matter health, both directly and indirectly via cardiovascular health. These results highlight the importance of cardiovascular risk factors for white matter health across the adult lifespan and suggest that systolic blood pressure, diastolic blood pressure and heart rate affect white matter via separate mechanisms.

## 1. Introduction

Cardiovascular function changes over the lifespan, with increasing risk of elevated blood pressure and heart rate in middle- and old age, and a lifetime risk for hypertension of 90% (Vasan et al., 2002). Such declines in cardiovascular health are also major risk-factors for clinically-silent brain injuries such as white matter lesions (including the white matter hyperintensities evident on magnetic resonance imaging scans; Iadecola and Davisson, 2008; Maniega et al., 2015; Verhaaren et al., 2013). White matter lesions, in turn, are associated with cognitive impairment and the risk of dementia, starting as early as the fifth decade (Knopman et al., 2001; Prins and Scheltens, 2015). This raises the possibility that the high prevalence of cardiovascular ill-health in middle- and old age contributes to the decline in cognitive functioning during later life (Kievit et al., 2016; Price et al., 2017; Prins and Scheltens, 2015). Unlike other risk factors of dementia such as variants of the *APOE* gene (Erikson et al., 2016), hypertension and elevated resting heart rate are preventable or potentially modifiable, by medication and by addressing behavioural risk-factors such as high body mass and lack of exercise (Gustafson et al., 2004; Ronan et al., 2016; Strömmer et al., submitted; Torres et al., 2015; Xu et al., 2013). At present, however, the relationship between cardiovascular and white matter health remains poorly understood, with most studies relying on a single indicator of white matter and cardiovascular health. To address this issue, we here used multivariate techniques to investigate the association multiple measures of cardiovascular health and white matter macro- and microstructure.

Most studies of cardiovascular health have investigated either hypertensive status or heart rate, but seldom both. There is cross-sectional and longitudinal evidence linking high systolic blood pressure to the development of white matter lesions (van Dijk et al., 2004; Verhaaren et al., 2013). For diastolic blood pressure, the evidence is inconclusive, with some studies finding that higher diastolic blood pressure predicts white matter lesions (Guo et al., 2009), while others find that lower diastolic blood pressure is associated with neurological abnormalities (Jochemsen et al., 2013). Elevated heart rate is also associated with cardiovascular problems, independent of blood pressure and physical activity (Fox et al., 2007; Woodward et al., 2012). However, few studies have directly investigated the association between heart rate and neurological disorders (e.g. Yamaguchi et al., 2015) and little is known about the relationship between heart rate and white matter macrostructure.

White matter degeneration is a continuous process, with microstructural changes occurring well before macrostructural changes (Maillard et al., 2013, 2014). Diffuse changes in white matter microstructure can be assessed using Diffusion Weighted Imaging (DWI), which yields measures of Fractional Anisotropy (FA), Mean Diffusivity (MD) and Mean Kurtosis (MK). While none of these measures have a clear one-to-one mapping to different histopathological processes (like myelination or axonal density), they show exquisite sensitivity to white matter microstructure (Jones et al., 2013; Maniega et al., 2015).

Most DWI studies on cardiovascular health have used FA, which measures the directionality of water diffusion in the brain (Jones et al., 2013). These studies have linked hypertension to decreased FA, likely indicating poor microstructural integrity (Hoptman et al., 2009; Maillard et al., 2012; Pfefferbaum et al., 2000). Notably, these microstructural changes may be particularly pronounced in frontal areas, modifiable by lifestyle factors such as Body Mass Index (BMI) and exercise, and predictive of cognitive ageing (Maillard et al., 2012; Raz et al., 2003; Strömmer et al., submitted).

Relatively few studies have investigated the effect of cardiovascular health on MD and MK (Maniega et al., 2015; Shimoji et al., 2014). However, these measures may provide complimentary information to FA. MD is particularly sensitive to vascular-driven white matter changes and correctly identifies up to 95% of white matter lesions, outperforming FA (Maniega et al., 2015). MK, unlike FA and MD, is sensitive to tissue microstructure and diffusion restrictions in regions with crossing or fanning fibres, and can therefore aid the assessment of brain-wide local tissue microstructure (Fieremans et al., 2011; Neto Henriques et al., 2015).

This review of the literature highlights that most studies to-date have relied on single indicators of both white matter and cardiovascular health, and employed mainly univariate statistical techniques, yielding fragmentary and partly inconsistent results, thus hampering translation into clinical practice. To systematically address this issue, we here used Structural Equation Modelling (SEM), a technique ideally suited to examining complex, multivariate relationships (Goldberger, 1972; McArdle, 2008).

SEM combines the strengths of path modelling (an extension of multiple regression) and latent variable modelling (McArdle, 2008). This feature of SEM allowed us to assess the relationship between multiple indicators of cardiovascular health (systolic blood pressure, diastolic blood pressure and heart rate), white matter macrostructure (white matter lesion volume and number), microstructure (FA, MD and MK in 10 white matter tracts) and lifestyle factors (BMI and exercise) in an adult lifespan sample of 667 participants aged 18 to 88 years.

We hypothesised that: (I) Cardiovascular health is associated with white matter lesion burden and microstructure (Guo et al., 2009; Maillard et al., 2012); (II) Systolic blood pressure, diastolic blood pressure and heart rate are each linked to white matter macro- and microstructure (Guo et al., 2009; Maillard et al., 2012; van Dijk et al., 2004; Yamaguchi et al., 2015); (III) BMI and exercise are associated with both white matter and cardiovascular health (Gustafson et al., 2004; Torres et al., 2015).

## 2. Methods

### 2.1. Sample

A healthy, population-based, adult lifespan sample was collected as part of the Cambridge Centre for Ageing and Neuroscience (Cam-CAN) study, as described in detail elsewhere (Shafto et al., 2014; Taylor et al., 2017). Exclusion criteria included low Mini Mental State Exam (MMSE ≤ 25), poor English, poor vision, poor hearing, MRI or MEG contraindications (for example, ferromagnetic metallic implants, pacemakers or recent surgery), self-reported substance abuse and other current serious health conditions (e.g. major psychiatric conditions). The final sample consisted of *N* = 667 cognitively healthy adults (*N*_cardiovascular measures_ = 579, *N*_white matter microstructure_ = 646, *N*_white matter macrostructure_ = 272, 50.60% female). See Table 1 for demographic information. Raw data can be requested from https://camcan-archive.mrc-cbu.cam.ac.uk/dataaccess/ and the covariance matrix and analyses scripts can be downloaded from https://github.com/dfl234/Cam-CAN Cardio White Matter Health.

**Table 1.**
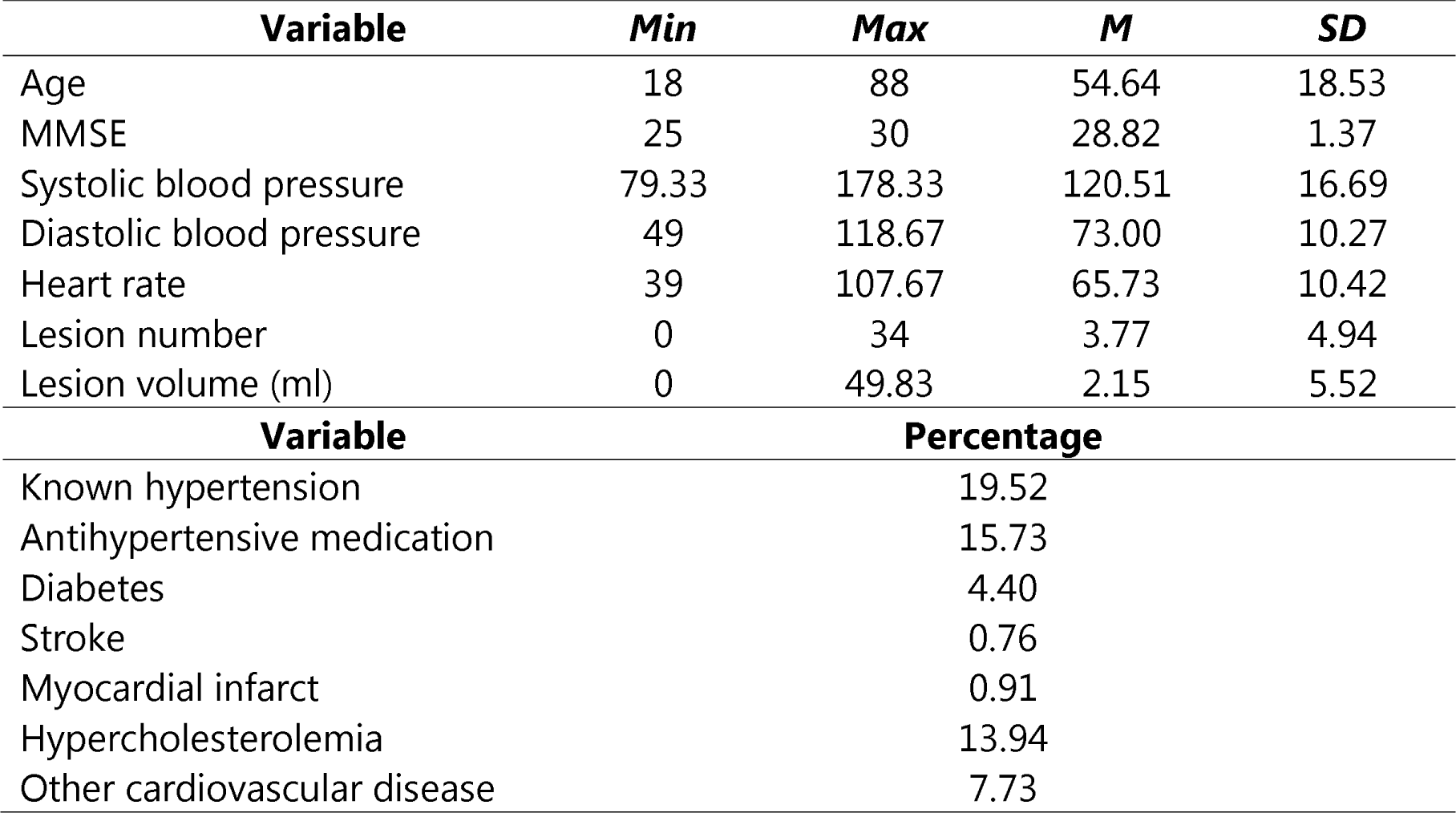
Demographic Information.

| **Variable** | **Min** | **Max** | **M** | **SD** |
| --- | --- | --- | --- | --- |
| Age | 18 | 88 | 54.64 | 18.53 |
| MMSE | 25 | 30 | 28.82 | 1.37 |
| Systolic blood pressure | 79.33 | 178.33 | 120.51 | 16.69 |
| Diastolic blood pressure | 49 | 118.67 | 73.00 | 10.27 |
| Heart rate | 39 | 107.67 | 65.73 | 10.42 |
| Lesion number | 0 | 34 | 3.77 | 4.94 |
| Lesion volume (ml) | 0 | 49.83 | 2.15 | 5.52 |
| **Variable** | **Percentage** |  |  |  |
| Known hypertension | 19.52 |  |  |  |
| Antihypertensive medication | 15.73 |  |  |  |
| Diabetes | 4.40 |  |  |  |
| Stroke | 0.76 |  |  |  |
| Myocardial infarct | 0.91 |  |  |  |
| Hypercholesterolemia | 13.94 |  |  |  |
| Other cardiovascular disease | 7.73 |  |  |  |

### 2.2. Cardiovascular Health

Heart rate, systolic and diastolic blood pressure were measured using the A&D Medical Digital Blood Pressure Monitor (UA-774), an automated sphygmomanometer. Measurements were carried out while participants were seated, and the arm resting comfortably. Blood pressure and heart rate are known to undergo fluctuation in response to stress and movement (Parati, 2005). We addressed this issue in two ways: First, to decrease the likelihood of confounding by recent movement, measurements were taken no sooner than 10 minutes after participants had been seated, and repeated three times in succession with approximately a one minute gap between measures. Second, latent variable scores were estimated from each of these measurements (see Structural Equation Modelling). Estimating latent variables from each measurement score can reduce measurement error and estimation bias and increase precision, compared to parcelling the data (e.g. averaging over measurement occasions) (Little et al., 2002, 1999).

### 2.3. White Matter Lesion Burden

White matter lesion volume and number were estimated automatically using the lesion growth algorithm provided by the LST toolbox for SPM (Schmidt et al., 2012): www.statistical-modelling.de/lst.html (Figure 1). This algorithm segments T2 hyperintense lesions using T1 and FLAIR images (see Supplementary Material for details of MRI data acquisition). Note that FLAIR images were collected as part of in-depth neurocognitive testing of a subsample of Cam-CAN participants, so that lesion burden was estimated for 272 participants only (Shafto et al., 2014). LST is a reliable, open-source, toolbox that demonstrates good agreement with manual tracing methods and very good specificity (Gracía-Lorenzo et al., 2013; Schmidt et al., 2012). The algorithm also meets general standards for automated lesion detection such as the use of both multimodal and spatial information (Gracía-Lorenzo et al., 2013). The algorithm requires user-specification of a threshold for transformed intensities *κ*. As recommended by Schmidt et al. (2012) the threshold intensity parameters *k* = 0.7 was chosen by visually inspecting lesion maps for different *κ* in four participants prior to modelling the data. We ran supplementary analyses with other thresholds *κ* to assess whether this choice impacted our results. We note that although the direction of effects and effect sizes for lesion volume were consistent across *κ*, the effect sizes for lesion number were lower for *κ* = 0.1 and *κ* = 0.3 (Supplementary Table 1).

**Figure 1.**
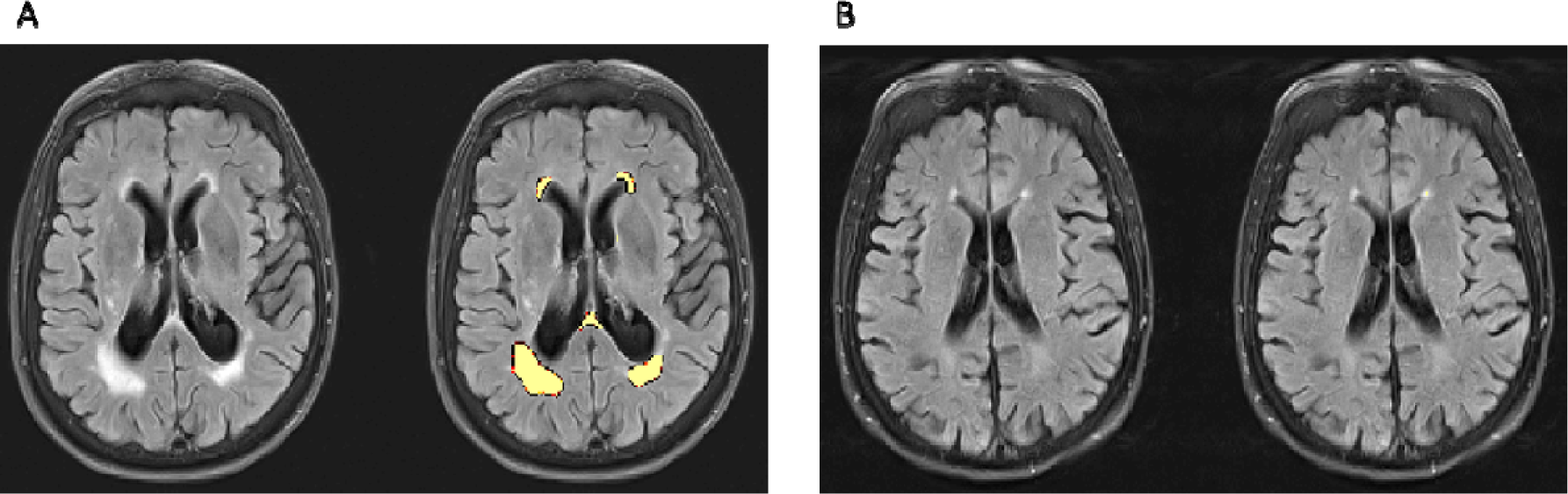
Estimating Lesion Burden. Panel A shows a participant with an estimated lesion number of 11 and volume of 29.16 ml. Panel B shows an age- and sex-matched control participant with a lesion number of 6 and volume of 0.72 ml. The left image in each panel shows T1-weighted images only, the right image shows T1-weighted images with lesion maps overlaid. Images were obtained using the LST toolbox for SPM (Schmidt et al., 2012).

### 2.4. White Matter Microstructure

To assess the relationship between cardiovascular health and white matter microstructure we modelled mean FA, MD and MK for ten tracts of the Johns Hopkins University (JHU) white matter tractography atlas averaged over the hemispheres (Figure 2). See Supplementary Methods for details of MRI data acquisition and pre-processing.

**Figure 2.**
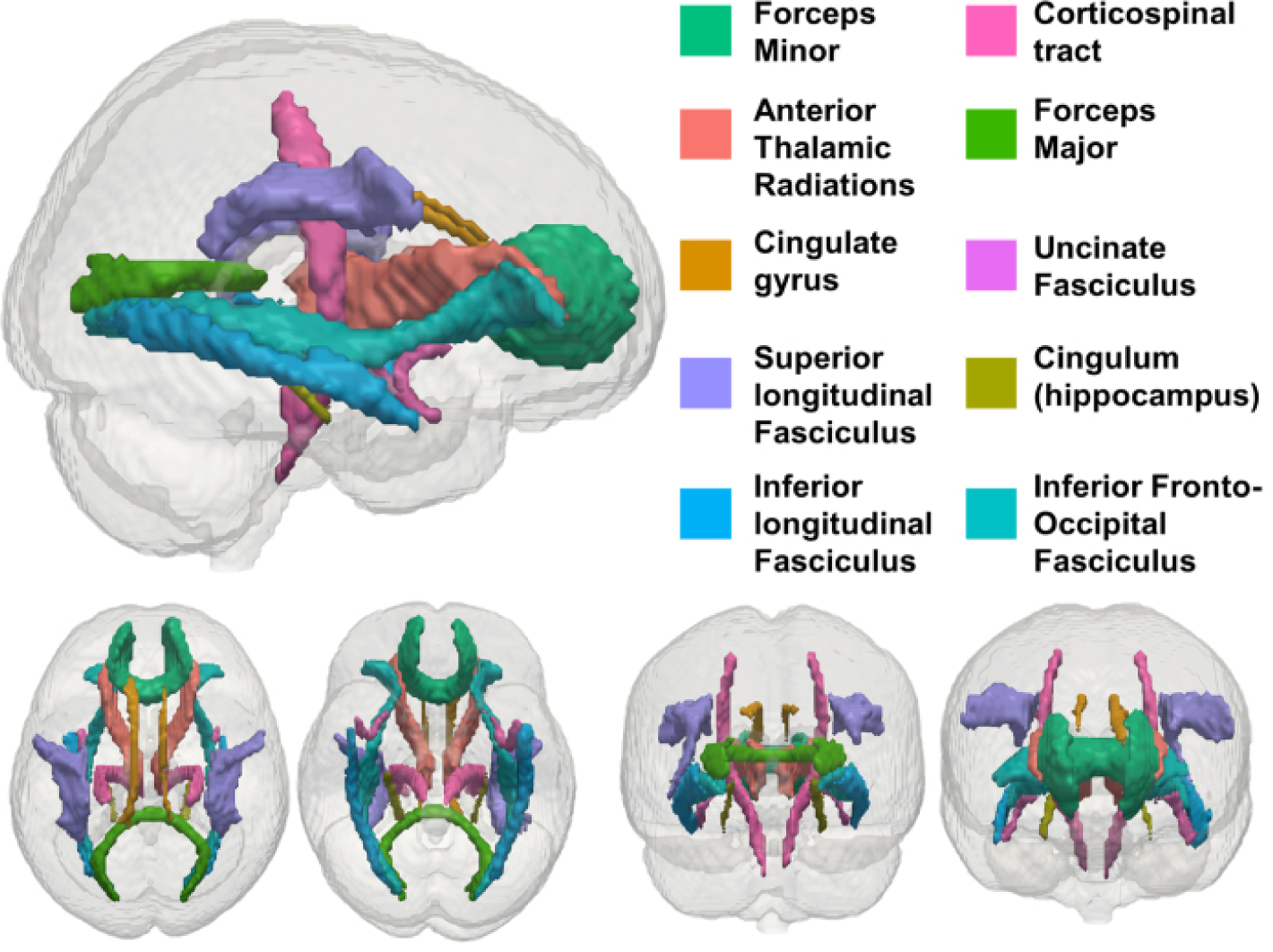
JHU White Matter Tracts Modelled in our Analysis. Adapted from (Kievit et al., 2016).

### 2.5. Protective Factors: Exercise and Body Mass Index

We modelled exercise and BMI as potential protective factors for cardiovascular and white matter health in an exploratory analysis. We assessed exercise using the European Physical Activity Questionnaire (Wareham et al., 2002). Four measures of physical activity energy expenditure in kJ/day/kg were calculated from self-reported physical activities at home, during leisure, at work and during the commute. Both paid employment and regular volunteering in the last 12 months was classified as work. BMI was calculated as weight (kg) / height (m)^2^. Height and weight were measured using portable scales (Seca 875).

### 2.6. Structural Equation Modelling

We modelled the relationship between cardiovascular health and white matter using Confirmatory Factor Analysis and SEM in R’s (R core team, 2015) Lavaan package (Rosseel, 2012). All models were fit using maximum likelihood estimation with robust (Huber-White) standard errors and a scaled test statistic. The residual variance of all observed variables was freely estimated. All available data was used. Missing data was estimated using the full information maximum likelihood method for all models. This method yields unbiased estimates when data is missing at random or missing completely at random, and as such is preferable to alternative techniques such as complete-case analyses or imputation methods (Enders & Bandalos, 2001). Model fit was inspected using the chi-square test, the Root Mean Square Error of Approximation (RMSEA) and its confidence interval, the Comparative Fit Index (CFI) and the Standardized Root Mean Square Residual (SRMR). We report the scaled test statistics. Good fit was defined as approximately RMSEA < 0.05, CFI > 0.97 and SRMR < 0.05, acceptable fit as approximately RMSEA = 0.08 − 0.05, CFI = 0. 95 − 0.97, SRMR = 0.05 − 0.10 (Schermelleh-Engel et al., 2003). Nested models were compared using a chi-square test. Effect sizes were evaluated by inspecting R2 for cardiovascular health and age overall and standardized parameter estimates for the individual effects of blood pressure and heart rate. Absolute estimates above 0.10 were defined as small effects, 0.20 as typical and 0.30 as large (Gignac and Szodorai, 2016).

## 3. Results

### 3.1. Measurement Models

In order to establish the relationship between cardiovascular and white matter health, we first used Confirmatory Factor Analysis to specify a measurement model for our cardiovascular health measures (Figure 3). Estimating latent variables has two benefits: First, it reduces measurement error in estimates of cardiovascular health (Little et al., 1999). Second, it allows for examining whether cardiovascular health is best represented as a single factor, which is theoretically plausible, or multiple, partially independent factors (here systolic blood pressure, diastolic blood pressure and heart rate). We first fit a single factor model, which represents cardiovascular health as a single latent dimension. This model did not fit well *χ*^2^(27) = 984.28, *p* < .001; RMSEA = .247 [.240 .255]; CFI = .474; SRMR = .218. We found that a three-factor model with heart rate, diastolic and systolic blood pressure as separate factors showed adequate fit (Figure 3; Supplementary Table 2), and fit better than the single-factor model (Supplementary Table 3; *χ*^2^(3) = 181.71, *p* < .001). We therefore used the three-factor measurement model in all subsequent analyses. The first measurement for both diastolic and systolic blood pressure showed a lower factor loading than the latter two measurements. This is consistent with the notion that the first measurement was less reliable, potentially due to movement or adaptation to the testing setting, and highlights the value of latent variables for reducing measurement error.

**Figure 3.**
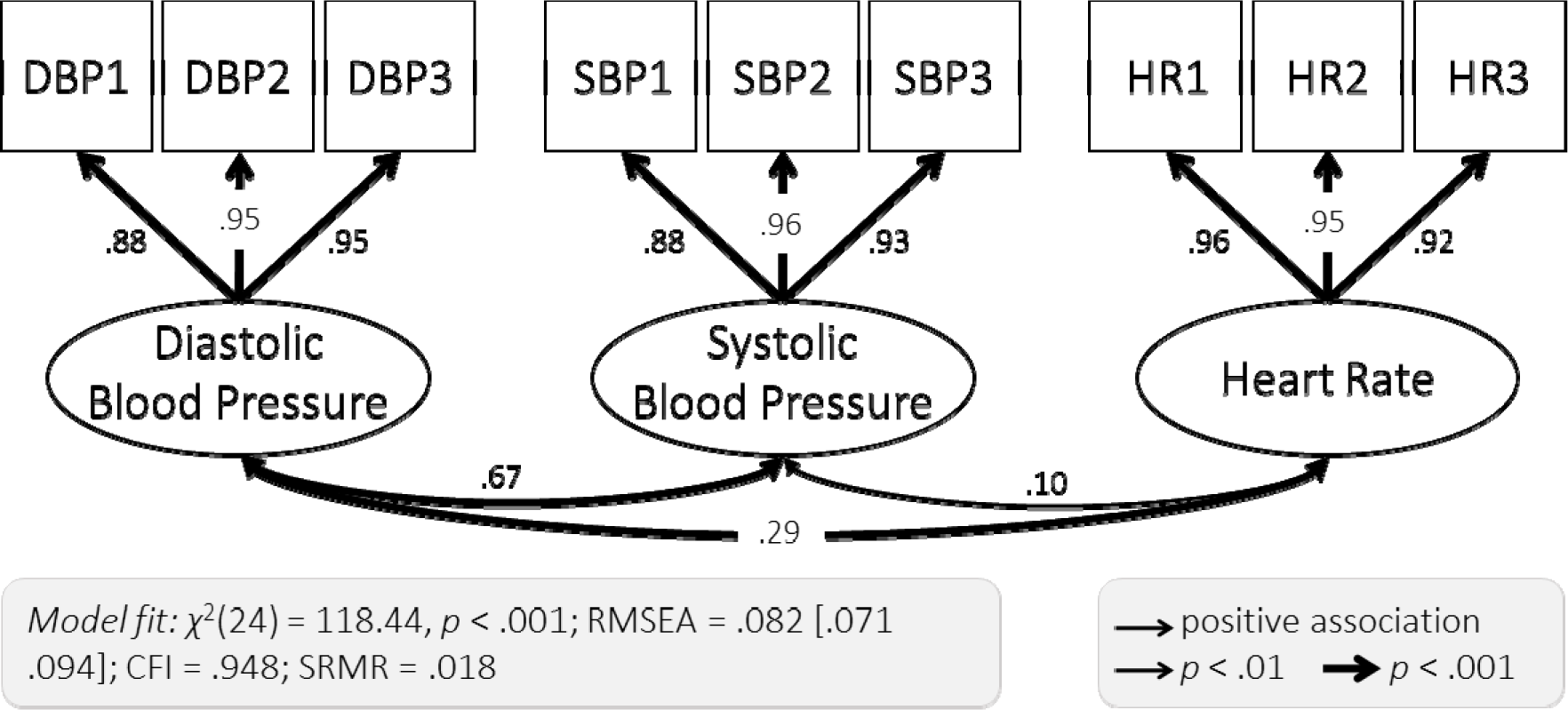
Three-Factor Measurement Model of Cardiovascular Health. Each of the three cardiovascular health factors was extracted from three measurements (diastolic blood pressure: DBP1, DBP2, DBP3, systolic blood pressure: SBP1, SBP2, SBP3 and heart rate: HR1, HR2, HR3). Standardized parameter estimates are shown.

We also assessed fit of a single-factor measurement model of white matter microstructure, by estimating a single latent variable from white matter microstructure in ten white matter tracts. We found that this single-factor model did not fit well for FA (*χ*^2^(35) = 418.66, *p* < .001; RMSEA = .130 [.120 - .140]; CFI = .879; SRMR = .062), MD (*χ*^2^(35) = 1825.69, *p* < .001; RMSEA = .281 [.271 - .292]; CFI = .766; SRMR = .090) or MK (*χ*^2^(35) = 283.51, *p* < .001; RMSEA = .105 [.097 - .112]; CFI = .840; SRMR = .054). This indicates that white matter microstructure cannot be adequately captured by a single factor in our cohort. We therefore modelled each of the ten white matter tracts as a separate indicator.

We were not able to assess model fit of a single-factor model of white matter lesion burden. There were only two indicators of lesion burden (total lesion volume and number), making a single-factor model just-identified. Model fit of just-identified models cannot be assessed. We therefore modelled each of the lesion burden measures as separate indicators in all subsequent analyses.

### 3.2. White Matter Lesion Burden

We examined the relationship between cardiovascular health and white matter lesion burden using Structural Equation Models in which diastolic blood pressure, systolic blood pressure and heart rate were modelled as predictors for total lesion volume and number. To account for potential confounding with age, this variable was additionally included as a covariate. The full model showed good fit (Figure 4). All regression paths, apart from the relationship between systolic blood pressure and lesion number, were significant (Figure 4), indicating that each cardiovascular measure made partially independent contributions to white matter lesion burden. Lower diastolic blood pressure, higher systolic blood pressure and heart rate each predicted greater total lesion volume and a higher number of lesions above and beyond age (Figure 4, Supplementary Table 1).

**Figure 4.**
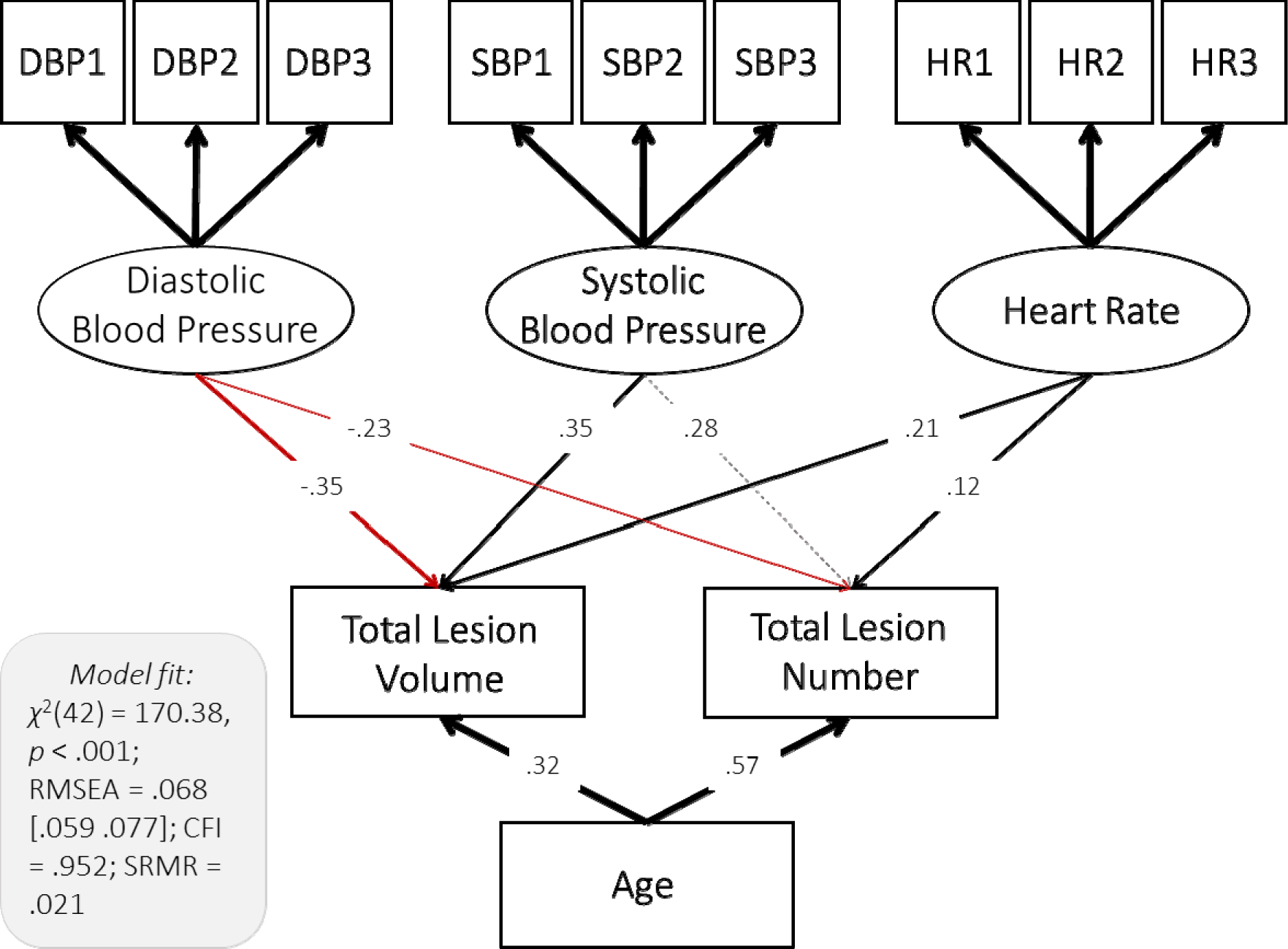
Path Model of the Relationship between Cardiovascular Health White Matter Lesion Burden. Diastolic blood pressure, systolic blood pressure and heart rate were modelled as latent variables (Figure 3). Age and total lesion volume and number were modelled as manifest variables. Standardized parameter estimates are shown. Residual covariances between cardiovascular factors, lesion burden measures and age were allowed but are not shown for simplicity.

Next, we examined whether diastolic blood pressure, systolic blood pressure and heart rate each had a specific link to white matter health, by comparing the freely-estimated model shown in Figure 4 to a model in which the parameter estimates for paths between cardiovascular health and each of the lesion burden measures are constrained to be equal (e.g. paths between lesion number and systolic blood pressure, diastolic blood pressure and heart rate). We found that the constrained model fitted worse than the freely-estimated model (*χ*^2^(4) = 31.49, *p* < .001). This indicates that diastolic blood pressure, systolic blood pressure, and heart rate differed in their relationship to white matter lesion burden, with diastolic and systolic blood pressure showing greater effects than heart rate (for standardized parameter estimates see Figure 4).

We took the same approach to testing whether lesion volume and lesion number showed different sensitivity to cardiovascular health. We found that a model in which parameter estimates for paths from each cardiovascular health measure to total lesion volume and number (e.g. paths between diastolic blood pressure and lesion volume and number) were constrained to be equal did not differ significantly in fit from a model in which these parameters were freely estimated (*χ*^2^(3) = 2.40, *p* = .494). This indicates that lesion volume and number showed a similar sensitivity to cardiovascular health. The total effect size of cardiovascular health and age on white matter lesion burden was considerable, *R*^2^ = 0.51 for lesion number and *R*^2^ = 0.30 for volume.

To better understand the negative association between diastolic blood pressure and lesion burden, we re-ran the model and included diastolic blood pressure as the only exogenous variable. We found that the association between diastolic blood pressure and total lesion volume and number became non-significant in this model (Supplementary Table 4). This indicates that the effects of diastolic blood pressure are conditional on other cardiovascular factors and age. This finding is compatible
with the notion that pulse pressure, the difference between systolic and diastolic
pressure, is a sensitive indicator of cardiovascular health (Kim et al., 2011; Strandberg and Pitkala, 2003). Exploratory models including pulse pressure and either systolic or diastolic blood pressure showed that higher pulse pressure was related to higher lesion burden overall (standardized coefficients ranging from 0.20 to 0.39, Supplementary Tables 5 and 6). Both systolic and diastolic blood pressure remained significant predictors of lesion burden over and above pulse pressure for lesion volume but not number (Supplementary Tables 5 and 6). This indicates that while pulse pressure is a sensitive indicator, absolute diastolic and systolic blood pressure can provide additional information about white matter health.

### 3.3. White Matter Microstructure

Next, we examined the link between cardiovascular health and non-clinical metrics of white matter microstructure. Specifically, we tested the relationship between cardiovascular health and FA, MD and MK in three separate models. For each of these models, diastolic blood pressure, systolic blood pressure, and heart rate were modelled as simultaneous exogenous variable. Age was included as a covariate. We found largely converging results across FA, MD and MK. For all models, blood pressure and heart rate made partially independent contributions to white matter microstructure. Consistently, lower diastolic blood pressure, higher systolic blood pressure, higher heart rate were each associated with lower FA, higher MD and lower MK (Figure 5, Table 2, Supplementary Figures 1 and 2). Out of our three measures of white matter microstructure, MD was generally most sensitive to cardiovascular health (Supplementary Table 7). Cardiovascular health and age explained up to 56% of the variance in MD (Supplementary Table 7), with absolute standardized parameter estimates indicating small to large effect sizes of cardiovascular health above and beyond age (Table 2). Effect sizes were generally larger for blood pressure than heart rate (Table 2). Although DWI metrics are only indirect measures of white matter microstructure (Jones et al., 2013), the associations reported here are compatible with an interpretation of adverse associations between cardiovascular ill-health and white matter microstructure (Falangola et al., 2008; Jones et al., 2013; Madden et al., 2012).

**Figure 5.**
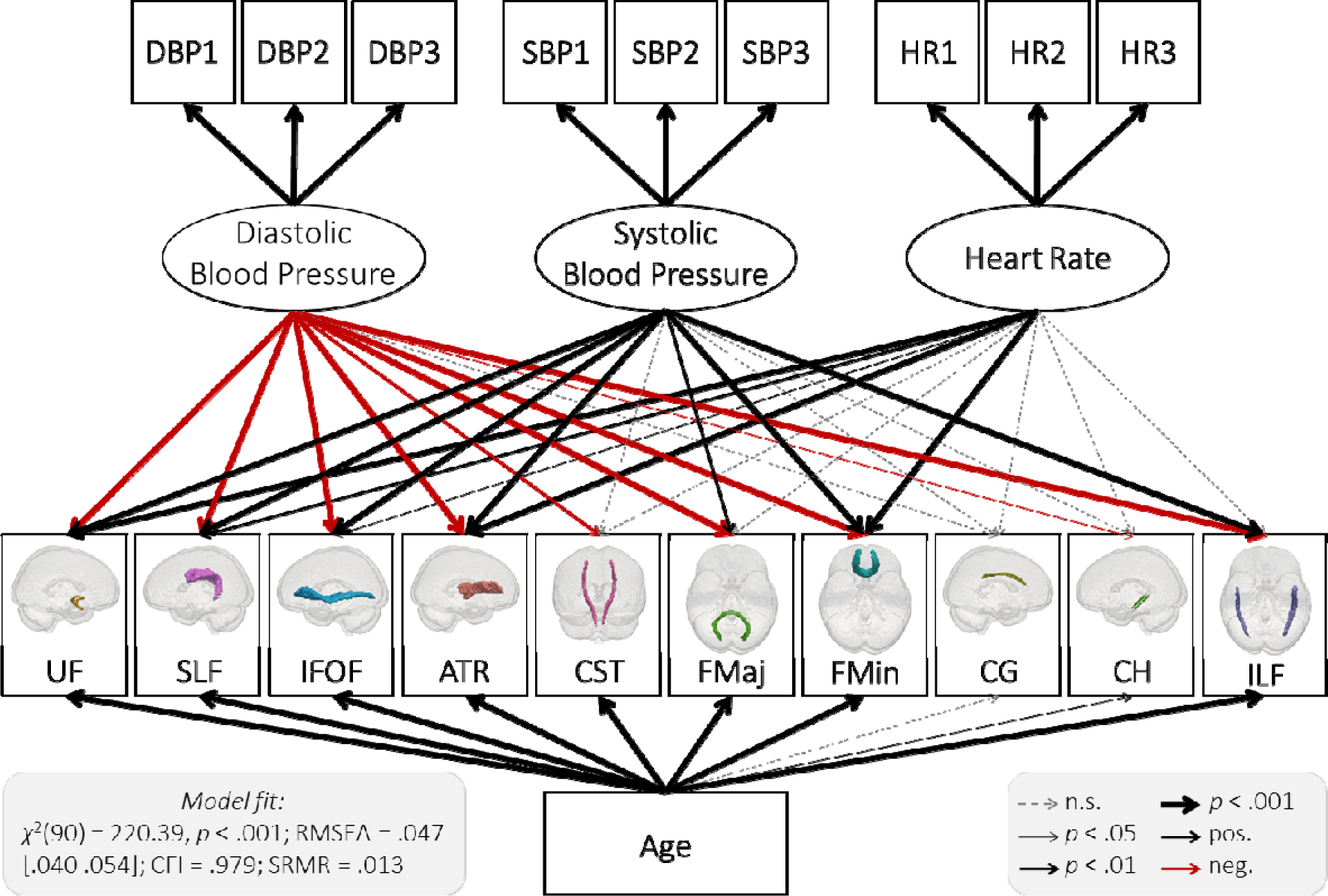
Path Model of the Relationship between Cardiovascular Health and MD. Diastolic blood pressure, systolic blood pressure and heart rate were modelled as latent variables (Figure 3). Age and MD in 10 white matter tracts were modelled as manifest variables. See Supplementary Table 9 for parameter estimates. Residual covariances between cardiovascular factors, lesion burden measures and age were allowed but are not shown for simplicity. Abbreviations: uncinate fasciculus (UNC), superior longitudinal fasciculus (SLF), inferior fronto-occipital fasciculus (IFOF), anterior thalamic radiations (ATR), cerebrospinal tract (CST), forceps major (FMaj), forceps minor (FMin), dorsal cingulate gyrus (CG), ventral cingulate gyrus (CH), inferior longitudinal fasciculus (ILF).

**Table 2.** Standardized Parameter Estimates for the Three Tracts Most Affected for FA, MD and MK.

|  | **FA** |  |  |  |
| --- | --- | --- | --- | --- |
|  | *DBP* | *SBP* | *HR* | *age* |
| Uncinate fasciculus | 0.19 | -0.26 | -0.15 | -0.23 |
| Anterior thalamic radiation | 0.18 | -0.15 | -0.15 | -0.40 |
| Inferior fronto-occipital fasciculus | 0.14 | -0.20 | -0.09 | -0.46 |
|  | **MD** |  |  |  |
|  | *DBP* | *SBP* | *HR* | *age* |
| Uncinate fasciculus | -0.36 | 0.38 | 0.17 | 0.38 |
| Superior Longitudinal Fasciculus | -0.30 | 0.26 | 0.11 | 0.55 |
| Forceps Minor | -0.25 | 0.23 | 0.12 | 0.62 |
|  | **MK** |  |  |  |
|  | *DBP* | *SBP* | *HR* | *age* |
| Uncinate fasciculus | 0.38 | -0.32 | -0.21 | 0.11 |
| Inferior fronto-occipital fasciculus | 0.36 | -0.32 | -0.14 | -0.32 |
| Forceps Minor | 0.35 | -0.30 | -0.17 | -0.34 |

*Note.* See Supplementary Table 8 – 10 for parameter estimates for all tracts.

To probe the specificity of the association between cardiovascular health and white matter microstructure, we again compared a series models with different equality constraints. For all measures of white matter microstructure, the full model fitted better than a model in which the parameter estimates of all cardiovascular factors were constrained to be equal (FA: *χ*^2^(20) = 49.53, *p* < .001; MD: *χ*^2^(20) = 118.66, *p* < .001; MK: *χ*^2^(20) = 53.16, *p* < .001). This highlights that diastolic blood pressure, systolic blood pressure, and heart rate differed in their relationship to white matter microstructure.

In terms of tract specificity, we found that the freely estimated FA and MD (but not MK) models fit better than models in which white matter tracts were constrained to be equally vulnerable (FA: *χ*^2^(27) = 54.80, *p* = .001; MD: *χ*^2^(27) = 44.59, *p* = .018; MK: *χ*^2^(27) = 30.54, *p* = .290), indicating that for FA and MD, white matter tracts differed in their sensitivity to cardiovascular health. The tracts most affected across our DWI measures were the uncinate fasciculus, inferior fronto-occipital fasciculus and forceps minor (Table 2).

Similar to lesion burden, the negative association between diastolic blood pressure and white matter microstructure was conditional on the effects of other cardiovascular health factors and age (Supplementary Table 11) and exploratory models including pulse pressure (systolic minus diastolic blood pressure) and either systolic or diastolic blood pressure showed that higher pulse pressure was related to poorer white matter microstructure (standardized estimates ranging from 0.10 to 0.41, Supplementary Tables 12 and 13). However, both systolic and diastolic blood pressure remained significant predictors of white matter microstructure over and above pulse pressure for most tracts (Supplementary Tables 12 and 13).

### 3.4. Modelling Protective Factors

In an exploratory analysis, we examined whether two potential protective factors for cardiovascular health and white matter health - BMI and self-reported exercise -affected cardiovascular and white matter health. We modelled exercise and BMI in separate models and assessed their direct and indirect (via cardiovascular health) effects on MD, the indicator of white matter health in our analyses that was most sensitive to cardiovascular health (Supplementary Table 7). For both exercise and BMI, a full model containing both direct and indirect effects on MD fit best (exercise: *χ*^2^(12) = 47.65, *p* < .001; BMI: *χ*^2^(3) = 128.67, *p* < .001). This indicates that exercise and BMI were associated with white matter microstructure both directly and indirectly via cardiovascular health.

#### 3.4.1. Exercise

The full model including exercise showed acceptable fit overall (*χ*^2^(117) = 577.52, *p* < .001; RMSEA = .077 [.071 .083]; CFI = .941; SRMR = .163). Exercise at home and during leisure showed no clear association with either MD or cardiovascular health (Supplementary Table 14). Exercise during commute showed no effect on MD but was the most significant predictor of cardiovascular health. More exercise during the commute correlated with lower diastolic blood pressure (standardized coefficient = −0.17, *p* = .001), systolic blood pressure (standardized coefficient = −0.20, *p* < .001), and heart rate (standardized coefficient = −0.13, *p* = .001), indicating that more energy spent while commuting was associated with better cardiovascular health overall. Exercise at work showed no clear relationship to cardiovascular health but was the most significant predictor of MD. There was a weak, negative correlation between exercise and MD in seven tracts with standardized coefficients ranging from −0.04 to −0.12 (Supplementary Table 14). This indicates that more exercise at work was associated with lower MD in these tracts, likely indicating better white matter health.

#### 3.4.2. Body Mass Index

The model including BMI showed mediocre fit overall *χ*^2^(102) = 693.70, *p* < .001; RMSEA = .093 [.087 .100]; CFI = .911; SRMR = .132. Higher BMI was associated with higher diastolic blood pressure (standardized coefficient = 0.29, *p* < .001), systolic blood pressure (standardized coefficient = 0.23, *p* < .001), and heart rate (standardized coefficient = 0.17, *p* < .001), thus predicting reduced cardiovascular health overall. Higher BMI was weakly correlated with lower MD in three tracts (Supplementary Table 15). This apparent negative effect of BMI was likely due to suppression. In a model where BMI alone was regressed onto MD, all but one non-significant association became positive. Here, lower BMI was significantly associated with lower MD, and therefore increased white matter microstructure, in four tracts (forceps minor, superior longitudinal fasciculus, inferior fronto-occipital fasciculus and anterior thalamic radiation), with standardized coefficients ranging from 0.08 to 0.18 (Supplementary Table 16).

### 3.5. Testing for potential confounds

Using multi-group models, we carried out a series of supplementary analyses of white matter lesion burden and microstructure to examine whether our results could be explained by possible differences between sexes, participants taking or not taking antihypertensive mediation and with and without known cardiovascular risk-factors (such as diabetes, hypercholesterolemia, etc., see Table 1 for full list). Results were invariant across groups for all of these factors (see Supplementary Analyses for details). Finally, we investigated whether social and lifestyle factors confounded our results by rerunning our model and including covariates for social class, education levels, smoking and alcohol consumption. The inclusion of these variables did not meaningfully change the directionality or significance level of the effects (see Supplementary Analyses for details).

## 4. Discussion

Here we show a link between common clinical measures of cardiovascular health and imaging indices of white matter health, in terms of both macro- and microstructure observed using multi-modal MRI in a population-based sample of healthy aging adults. Lower diastolic blood pressure, higher systolic blood pressure and higher heart rate were each strongly, and independently associated with poorer white matter health on all indices - over and above the effects of age. The link between cardiovascular and white matter health was robust across genders, in people taking and not taking antihypertensive medication, with and without known cardiovascular risk-factors (diabetes, elevated cholesterol levels etc.), and when controlling for social class, education levels, alcohol consumption and smoking.

### Systolic hypertension and diastolic hypotension

We found that systolic hypertension predicted poorer white matter macro- and microstructure on all indices, in line with previous studies showing bivariate links between high systolic blood pressure and lower FA, higher MD, and greater white matter lesion volume and number (Maillard et al., 2012; van Dijk et al., 2004; Verhaaren et al., 2013). Systolic blood pressure is known to increase more steeply with age than diastolic blood pressure and has been argued to be a better predictor of cardiovascular and neurological outcomes (Strandberg and Pitkala, 2003). However, we find that *lower* diastolic blood pressure had a similarly detrimental effect as *higher* systolic blood pressure. This negative correlation was not evident in a single-indicator model including diastolic blood pressure alone, showing that low diastolic blood pressure is likely to be particularly detrimental at moderate to high levels of systolic blood pressure. This conditional effect may capture the difference between systolic and diastolic blood pressure, a measure known as *pulse pressure* (Kim et al., 2011; Strandberg and Pitkala, 2003). Pulse pressure is often argued to be a particularly good measure of cardiovascular health in older populations because systolic and diastolic blood pressure widen with age (Supplementary Figure 3), increasing pulse pressure, and contributing to the high prevalence of isolated systolic
hypertension in older adults (Huang et al., 2004; Strandberg and Pitkala, 2003). Exploratory models showed that pulse pressure was indeed a sensitive indicator of white matter micro- and macrostructure and showed greater effect sizes than either systolic or diastolic blood pressure on their own. However, both systolic and diastolic blood pressure remained significant predictors of white matter health after controlling for pulse pressure, indicating that they each capture information about cardiovascular health over and above pulse pressure. Pathophysiologically, isolated systolic hypertension in combination with diastolic hypotension may reflect reduced vessel compliance and increased arterial stiffness, one of the hallmarks of vascular ageing (Vlachopoulos et al., 2010).

Overall, these findings highlight the importance of assessing and modelling systolic and diastolic blood pressure, rather than just hypertensive status. The use of multivariate models reveals the existence of conditional effects which can remain undetected when using univariate techniques: lower diastolic blood pressure in combination with higher systolic blood pressure was particularly detrimental for white matter health. This pattern of results is also relevant to clinical practice. Current UK treatment guidelines include only upper limits for blood pressure (NICE, 2016), but our findings indicate that lower limits for diastolic blood pressure, or pulse pressure, may provide crucial complementary information about cardiovascular health and its consequences.

### Elevated heart rate

Elevated heart rate was associated with poorer white matter health independent of age and blood pressure. Previous research showed that high heart rate predicts cardiovascular problems independent of blood pressure, physical activity and comorbidities (Cooney et al., 2010; Fox et al., 2007; Woodward et al., 2012). Nighttime heart rate has also been implicated in white matter lesions and stroke (Yamaguchi et al., 2015). Here, we demonstrate that higher heart rate during the day is associated with poorer white matter microstructure and macrostructure, even though effect sizes were somewhat smaller than for blood pressure. The aetiology and impact of elevated heart rate remains poorly understood but may be related to sympathetic nervous system hyperactivity and endothelial dysfunction, which, in turn, may put stress of vascular architecture (Palatini, 2011).

### Body mass and exercise

Body mass and exercise were related to white matter health, both directly and indirectly via cardiovascular health. Higher BMI was associated with poorer cardiovascular health and higher MD, indicating reduced white matter integrity, even though effect sizes were relatively small. This finding replicates previous studies showing a negative association of BMI and white matter health (Gustafson et al., 2004; Ronan et al., 2016) and indicates that both cardiovascular and white matter health are likely modifiable by controlling BMI. For exercise, exercise at work and exercise during the commute were significant, albeit weak, predictors. Exercise at home and during leisure showed no clear effect. This dissociation may reflect the regularity of exercise: Exercise at work and during the commute is likely more regular than exercise at home and during leisure. It may also reflect factors that are not, or not solely, related to physical activity. The exercise questionnaire used here classed regular volunteering as work (Wareham et al., 2002). Therefore, exercise at work and during commute may reflect social and intellectual engagement and their protective effects - particularly during retirement (James et al., 2011). Future research will have to disentangle these potential mechanisms of cardiovascular and white matter health and include in-depth analyses of physical and mental health, as well as social and cognitive engagement, and their links to cardiovascular and brain health. This will help inform interventions to promote healthy cognitive ageing.

### Limitations

Despite strengths of this study, such as sample size, multivariate methodology and a broad set of cardiovascular and neural measures, several limitations should be noted. First, given the inclusion criteria for Cam-CAN imaging (see Taylor et al., 2017), our sample are likely *more* healthy than the average population. While this limits generalizability, it suggests that the negative effects of poor cardiovascular health are observed even in healthy, non-clinical samples.

Furthermore, blood pressure and heart rate are known to undergo fluctuation in response to stress and movement (Parati, 2005). Although we measured heart rate and blood pressure three times and modelled them as latent variables, reducing measurement error and increasing precision (Little et al., 1999), ambulatory, 24-h assessment of blood pressure and heart rate may be even more sensitive. Such methods may be able to pick up additional disease symptoms such as variability in blood pressure or a lack of nocturnal non-dipping of heart rate (Fox et al., 2007; Rothwell et al., 2010; Yamaguchi et al., 2015). Nonetheless, our findings indicate that even simple measurements of heart rate and blood pressure, as may be taken during a single health assessment, may have considerable prognostic value.

Similarly, our findings of the effects of exercise depend on limitations of questionnaire measures in accurately assessing levels of exercise (Prince et al., 2008). Future research may want to use direct assessments of physical activity, like accelerometers, which are generally more reliable than self-report measures (Prince et al., 2008). This may help tease apart the effects of exercise from other lifestyle factors such as social and intellectual engagement.

Finally, although previous longitudinal research in humans (Dufouil et al., 2001; Gustafson et al., 2004) and experimental work in animal models (Kaiser et al., 2014; Zuo et al., 2017) suggests causal links between cardiovascular and white matter health, our study was cross-sectional. As such, we cannot establish the temporal order of particular cardiovascular symptoms and changes in white matter micro- and macro- structure. We included age as a covariate in all structural equation models, thus accounting for possible linear age associations. Although non-linear associations can be estimated using structural equation models, we limit ourselves here to more parsimonious linear associations only. We cannot, however, completely rule out other alternative explanations such as cohort effects. Our findings should therefore ideally be interpreted alongside interventional and longitudinal studies to further establish causal links. Future experimental studies will also need to investigate possible disease pathways, which, at present, are only partially understood (Iadecola and Davisson, 2008).

### Implications for cognitive ageing

We found that the uncinate fasciculus, forceps minor and inferior fronto-occipito fasciculus were the three white matter tracts most affected by cardiovascular health. While we did not explicitly model cognitive function in the present study, we note that these three tracts have been consistently linked to cognitive ageing in previous research. The uncinate fasciculus is thought to play a major role in the formation of mnemonic associations and episodic memory (von der Heide et al., 2013; but see Henson et al., 2016) and has been implicated in the development of cognitive impairment in dementia (Serra et al., 2012). The forceps minor has been shown to predict processing speed and fluid intelligence in the Cam-CAN cohort (Kievit et al., 2016). The inferior fronto-occipital fasciculus has been linked to linked to linguistic processing (Dick and Tremblay, 2012) and set-shifting (Perry et al., 2009). This pattern of results is therefore consistent with the notion that cardio-vascular ill-health may contribute to cognitive decline during ageing.

## Conclusion

Our multivariate analyses show that lower diastolic blood pressure, higher systolic blood pressure and higher heart rate were each strongly associated with poorer white matter macro- and microstructure in a large sample of healthy, community dwelling adults. These adverse effects were largely additive – diastolic blood pressure, systolic blood pressure and heart rate each made partially independent contributions to white matter health. These findings highlight the value of concurrently assessing multiple indicators for cardiovascular health and indicate that systolic blood pressure, diastolic blood pressure and heart rate may each affect neurological health via separate disease mechanisms.

## Acknowledgements

The authors would like to thank Carol Brayne, Simon Davis and Rik Henson for helpful comments on the study design and imaging techniques. DF, DN, JBR, DP and RAK are supported by the UK Medical Research Council. The Cambridge Centre for Ageing and Neuroscience (Cam-CAN) was supported by the Biotechnology and Biological Sciences Research Council (grant number BB/H008217/1). RAK is supported by the Sir Henry Wellcome Trust (grant number 107392/Z/15/Z) and MRC Programme Grant MC-A060-5PR60. This project has also received funding from the European Union’s Horizon 2020 research and innovation programme (grant agreement number 732592).

The Cam-CAN corporate author consists of the project principal personnel: Lorraine K Tyler, Carol Brayne, Edward T Bullmore, Andrew C Calder, Rhodri Cusack, Tim Dalgleish, John Duncan, Richard N Henson, Fiona E Matthews, William D Marslen-Wilson, James B Rowe, Meredith A Shafto; Research Associates: Karen Campbell, Teresa Cheung, Simon Davis, Linda Geerligs, Rogier Kievit, Anna McCarrey, Abdur Mustafa, Darren Price, David Samu, Jason R Taylor, Matthias Treder, Kamen Tsvetanov, Janna van Belle, Nitin Williams; Research Assistants: Lauren Bates, Tina Emery, Sharon Erzinlioglu, Andrew Gadie, Sofia Gerbase, Stanimira Georgieva, Claire Hanley, Beth Parkin, David Troy; Affiliated Personnel: Tibor Auer, Marta Correia, Lu Gao, Emma Green, Rafael Henriques; Research Interviewers: Jodie Allen, Gillian Amery, Liana Amunts, Anne Barcroft, Amanda Castle, Cheryl Dias, Jonathan Dowrick, Melissa Fair, Hayley Fisher, Anna Goulding, Adarsh Grewal, Geoff Hale, Andrew Hilton, Frances Johnson, Patricia Johnston, Thea Kavanagh-Williamson, Magdalena Kwasniewska, Alison McMinn, Kim Norman, Jessica Penrose, Fiona Roby, Diane Rowland, John Sargeant, Maggie Squire, Beth Stevens, Aldabra Stoddart, Cheryl Stone, Tracy Thompson, Ozlem Yazlik; and administrative staff: Dan Barnes, Marie Dixon, Jaya Hillman, Joanne Mitchell, Laura Villis.

